# The influence of movement speed on reward-based motor learning

**DOI:** 10.1101/2023.06.28.546754

**Authors:** Nina M. van Mastrigt, Katinka van der Kooij, Jeroen B. J. Smeets

**Affiliations:** Vrije Universiteit Amsterdam, Department of Human Movement Sciences, Amsterdam, The Netherlands

## Abstract

Human movement is inevitably variable. This variability can be seen as a constraint to overcome, but it may also be a feature: being variable may result in the discovery of better movement solutions. Especially when feedback is limited to binary information on movement success or failure, variability is key for discovering which movements lead to success. Since moving faster increases variability, we aimed to answer the question whether movement speed can be harnessed to improve such reward-based motor learning. Subjects performed a stepping task in a slow and a fast session. They had to learn the gain between their step lengths and visual target distances on screen based on binary reward feedback. We successfully manipulated movement speed between sessions and participants could learn the gain in both sessions. We found no difference in learning between speed sessions, despite the fact that variability in gain increased in the fast relative to the slow session. To distinguish between different sources of variability, we estimated inevitable motor noise from the variability following successful trials. We estimated exploration as the additional variability following non-successful trials relative to following successful trials. We found no relation between variability sources and learning. In conclusion, reward-based motor learning is possible in a gain-learning task. In this task, moving faster did not lead to higher learning. Since the role of variability may differ between experimental tasks, whether movement speed can be harnessed to improve motor learning needs to be tested in other experimental tasks.

## Introduction

A well-known characteristic of human movement is that it is inevitably variable: humans cannot repeat a movement in exactly the same way (Bernstein, 1967). Another well-known characteristic of human movement is that faster movements are less accurate (Fitts, 1954), hence more variable. Variability in movement has long been thought to only act as a constraint to overcome (Harris & Wolpert, 1998): by limiting accuracy, it may hamper performance. A more recent view on motor variability is that it may also be a feature: by yielding variable information about the body and environment, it may result in the discovery of better movement solutions (Herzfeld & Shadmehr, 2014). This leads us to the question whether the trade-off between speed and accuracy can be harnessed to promote motor learning.

The idea that motor variability may serve as a feature stems from reinforcement learning theory (Sutton & Barto, 2017), which states that learning from binary information on success and failure requires exploration of which action leads to success. Since such exploration is impossible without variability in movements, variability is key to reinforcement learning. Indeed, variability is regulated during learning from binary reward feedback in song birds (Tumer & Brainard, 2007), rats (Dhawale et al., 2019) and humans (Abram et al., 2022; Cashaback et al., 2019; Pekny et al., 2015; Therrien et al., 2016, 2018; Uehara et al., 2019; van der Kooij et al., 2021; van der Kooij & Overvliet, 2016; van der Kooij & Smeets, 2018; van Mastrigt et al., 2020). Some evidence exists for a positive relationship between learning and baseline variability or regulation of variability (Dhawale et al., 2019; Wu et al., 2014), but this evidence is mainly correlational and not convincingly replicated (He et al., 2016).

The relation between motor variability and motor learning may depend on the contributions of different sources of motor variability. In error-based motor learning, motor variability is thought to result from both central planning noise and peripheral execution noise (van Beers, 2009; van der Vliet et al., 2018). In reward-based motor learning, motor variability is thought to result from inevitable sensorimotor noise and exploration in response to feedback (Cashaback et al., 2019; Dhawale et al., 2019; Therrien et al., 2016; van Mastrigt et al., 2020). Using a reward-based motor learning paradigm, (Therrien et al., 2018) found that smaller ratios of exploration to sensorimotor noise were associated with less learning. The relation between motor variability and motor learning likely depends on the contributions of different sources of variability. Whether the trade-off between speed and accuracy can be harnessed to promote learning might depend on the type of variability that is increased by moving faster.

Here, we aimed to investigate the relation between movement speed and reward-based motor learning from the perspective of motor variability. Subjects performed a stepping task with binary reward feedback at a slow and a fast movement speed. We first checked whether increased movement speeds lead to higher variability as prescribed by the speed-accuracy trade-off (Fitts, 1954). We then tested whether learning depended on speed to answer the question whether increasing movement speed can be harnessed to promote learning. We explored how movement speed influences different sources of variability and how (changes in) sensorimotor noise, exploration and their ratio are related to (changes in) learning.

## Methods

### Participants

After providing informed consent, forty adults (reported mean age 22 (+/-5 years), 8 male) participated in this study. Participants were recruited from a study participation pool for Bachelor students at the Vrije Universiteit Amsterdam as well as from the personal network. Ethical approval for the study was provided by the local ethical committee of the Vrije Universiteit Amsterdam.

### Set-up

Participants stood on a custom made 1×1 meter eight-sensor strain gauge force plate that recorded center-of-pressure position with a sampling frequency of 100 Hz (Figure 1). They stood 2 meters in front of a 54.7 × 31.5 cm screen (Asus LCD, 1920 × 1080 pixels) that depicted stimuli with a refresh rate of 240 Hz (Figure 1). Stimuli were generated with an experimental script programmed in Open Sesame (Mathôt et al., 2012) on a Windows 10 computer (Dell, Intel Core i7 vPro), while center-of-pressure data were collected with a custom made programme on another Windows 10 computer (HP precision 3630, Intel Core i7). Forward center-of-pressure position data of the last 600 ms of each movement were sent to the Open Sesame computer using a COM connection with a baud rate of 9600. Participants were prevented from falling off the force plate in the backward direction by a 4 cm aluminium bar at 0 and 1 m height (Figure 1).

**Figure 1.**
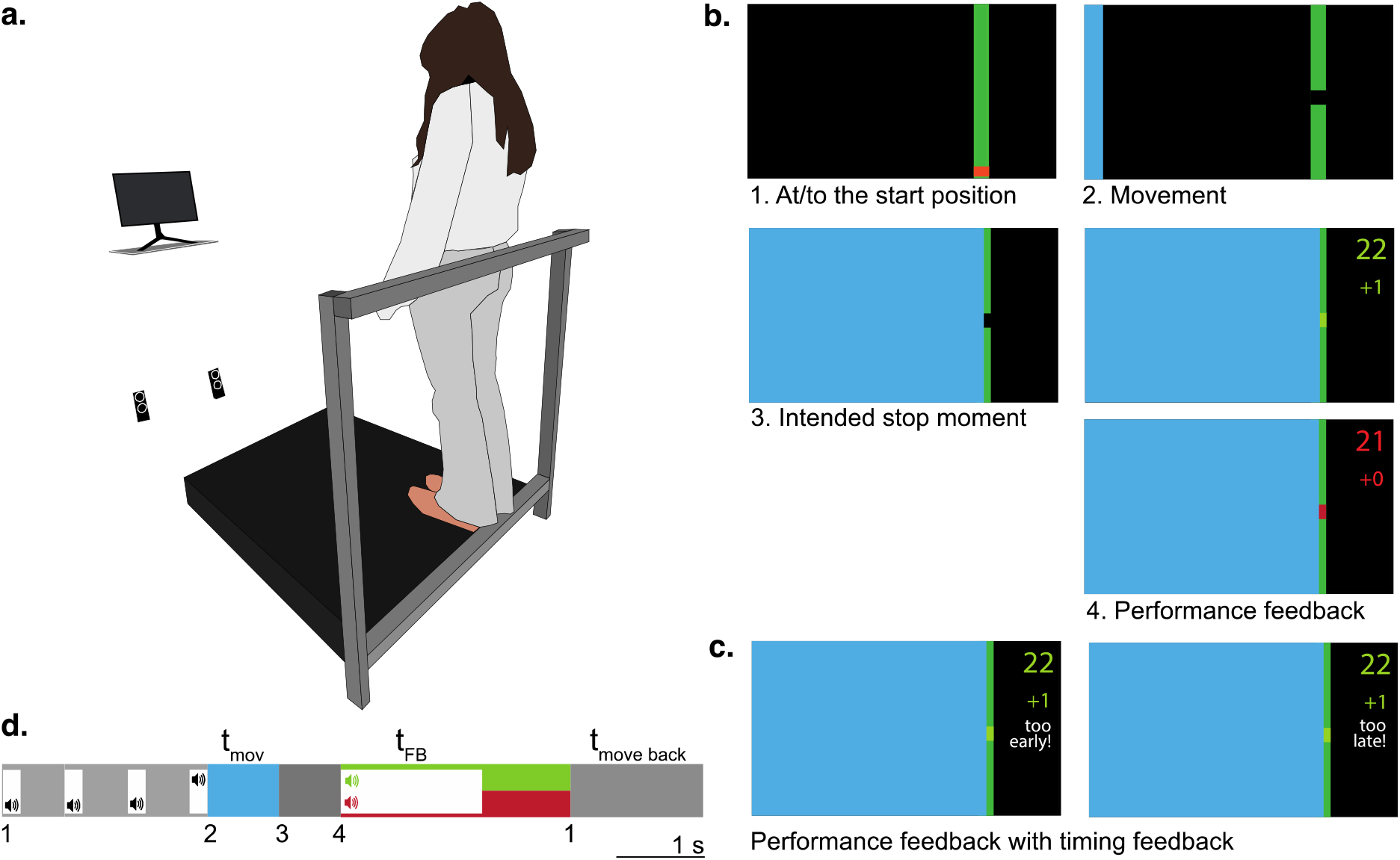
Stepping task. a. Set-up b. Sequence of visual stimuli of one trial, including performance feedback. 1. Participants start with their heels against a bar, corresponding to the orange square at the lower end of the green dam. 2. A gap appears and the blue see approaches. 3. Participants move their feet in the direction of the gap to block it when the see hits the dam. 4. Participants receive performance feedback on whether they hit (light green) or missed (red) the gap. c. When people stopped moving too early or too late, timing feedback was added to the performance feedback, independent of reward. d. Timeline of one trial. Numbers refer to screens in panel b.

### Task

Participants performed a goal-directed stepping task. On the screen, participants saw a vertical bar that divided the screen in two areas (Figure 1b). Participants were told this bar was a top-down view of a dam (green) protecting the country (right area) from flooding by the approaching sea (blue area on the left) (Figure 1b). Participants were instructed that when a gap appeared in the dam, they had to fill the gap by stepping forward with some unknown correct step length. Participants had to learn the gain, i.e. the relation between correct step length and distance to the gap. Participants received binary reward feedback based on their step length, i.e. the forward distance travelled in a step.

Every trial started from a fixed start position at the back of the force plate, where participants’ heels touched a bar (Figure 1a). Participants saw this start position as an orange box on the screen (Figure 1b). After four beeps of 200ms with 500 ms in between (Figure 1d), a new target would become visible as a black gap in the green dam. At the same time, the sea started flowing towards the dam with a constant velocity (Figure 1b, screen 2). Participants were instructed to start stepping at the fourth beep and to stand still with two feet side-by-side on the floor when the sea arrived at the middle of the gap (Figure 1b, screen 3). 700 ms after each attempt, participants received feedback on both performance and timing (Figure 1b-d). Reward feedback consisted of a 1.6 s “bing” sound, the gap turning green and a green +1 in case of success (Figure 1d), and a 1.6 s “teh-deh” sound, the gap turning red and a red +0 in case of failure (Figure 1d). A total score per block was shown as well (Figure 1b). If the timing of standing still was approximately correct, timing feedback consisted of no message; else “too early” or “too late” was shown in smaller white letters (Figure 1c). Participants viewed the feedback for 2.6 s and then had 1.5 s to step back to their start position (Figure 1d). After an inter-trial interval of 500 ms, the next trial started.

#### Experimental sessions: speed

Each participant performed two experimental sessions that differed in movement speed: a Slow and a Fast session. Movement speed (i.e. time to make the movement) was manipulated by the speed of the sea flowing from left to the middle of the gap. The intended movement duration was 800 ms (Slow) or 400 ms (Fast). As movement speed is distance divided by time, movement speed on each trial depended on the step length. The two sessions were performed directly after each other with a short break in between, in random order.

### Procedure

Participants started with a baseline starting position measurement in which they had to stand still for 50 seconds. They received instructions on the screen that their task would be to fix gaps in a dam on screen, by using the force plate as a tool. By stepping forward with two feet, they could control where the missing piece would end up. They were instructed that step length determined success, that there would be a new relation between the gap distance on the screen and the correct step length. To demonstrate what this relation was, they were shown a gap halfway the screen and were asked to take a step halfway the force plate. They were then shown where the missing piece ended up, and were asked to do another attempt to make the piece end up in the gap. Next, they were instructed to start moving at the fourth beep, and to come to stand still the moment the border of the blue sea arrives at the middle of the gap. They were told that they would receive points if they performed better than the previous 10 trials. Next, they performed two practice sessions. The first practice session was to train movement speed. It consisted of two blocks of 25 trials with a fixed gap distance and timing feedback only, first for slow trials and then fast trials. The second practice session was to practice with different gap distances and performance feedback. It consisted of two blocks of 5 trials with varying gap distances, a gain of 1 and both timing and performance feedback. After the practice, participants performed the slow and fast experimental session in random order. Each session consisted of five blocks of 50 trials; the gain was constant within a block but switched between blocks.

#### Gains

From block to block, participants had to learn a new gain. We defined the gain as the correct step length divided by the gap distance. Distances were expressed as a fraction of the size of the force plate and screen, respectively. The gains of the five blocks were always 1.4, 0.83, 1.2 and 0.67. Gains larger than 1 indicate that relative stepping distance was larger than relative visual distance. Gains smaller than 1 indicate that relative stepping distance was smaller than relative visual distance. The order of these blocks remained fixed, but the first gain was randomly determined with the constraint that the two speed sessions could not start with the same gain. The fifth block always had the same gain as the first block. This way, each gain was always preceded by a fixed other gain.

#### Targets

Gap positions were randomly generated using three constraints. A random position on the force plate was chosen that was constrained based on force plate length. Firstly, gap positions on the force plate were constrained so that participants never had to make steps to positions that fell within 5% and 95% of the force plate length. Secondly, they should never be required to make steps smaller than 25% of the force plate size and to make steps bigger than 75% of the force plate length. For this, step length was calculated by subtracting start position. Next, the force plate position was converted to a screen position by multiplying with the inverse gain of that block. Thirdly, gap positions on screen were constrained so that they fell within 36% and 74% of the screen height to prevent outliers in targets to reveal whether the gain was smaller or larger than 1.

#### Feedback

Feedback was based on the unfiltered center-of-pressure recorded by the force plate. The start position was defined as the median forward center-of-pressure position of ten 600-ms samples of a period of 50 seconds prior to the experiment, in which the participant stood still with the heels touching the bar at the back of the force plate. Step length was defined as the difference between the median forward centre-of-pressure position in the 400 ms after the intended stopping moment and the start position.

*Performance feedback* Success was determined based on gain according to a combination of a fixed and an adaptive reward criterion. In the first ten trials of each speed session, a fixed reward criterion defined gains within a fixed zone around the target gain as successful. The upper and lower border of this reward zone were determined by the correct relative step distance for the upper and lower border of the gap (i.e. relative screen distance times target gain), divided by the relative screen distance of the middle of the gap. For later trials, we used an adaptive reward criterion to make participants step both more accurately and less variably. The width of the reward zone around the target gain was based on the gains of the previous ten trials. For these previous ten trials, we calculated the median gain and based on this, the gain bias as 90% of the absolute error between this median gain and the target gain of that block. For the previous ten trials, we also calculated the variability as half of the interquartile range in the gains. If the gain bias was larger than the variability, the width of the reward zone was set to the gain bias. If the variability was larger the gain bias, the width of the reward zone was set to the variability. The adaptive reward criterion was used to make sure that people could find the target gain and to make sure that we would have equal numbers of rewarded and non-rewarded trials for estimating exploration.

*Timing feedback* Timing feedback was about arriving too early or too late. The feedback was based on forward center-of-pressure position in the 200ms before and 400 ms after the moment the sea arrives at the dam. In this interval, we started looking for the first moment that the participant stopped. We detected the first sample for which velocity (i.e. a two-sample difference) was negative and defined this as the stop moment. To prevent noise in the CoP signal from influencing the detection of the stop moment, we constrained stop detection to a final movement interval: the time interval between the time point with the maximum forward position during that trial and the time point at which 80% of this value was reached. After detecting the stop moment, we checked whether it was within a certain interval around the intended stopping moment. If it was, timing was judged to be correct. If the first stop sample was before or after that interval, timing feedback was provided (‘too early’ or ‘too late’, respectively). In the Slow session, the interval was 360 ms. In the fast speed session, the interval was 180 ms.

### Data analysis

#### Movement speed

For each trial, we calculated movement speed at each time point within the intended movement period by taking the derivative of the forward center-of-pressure position. We defined average movement speed as the average over all time points per trial. To obtain the average movement speed per gain block, we took the mean of all trials of each gain block.

To obtain one value for movement speed per speed session, we took for each participant the mean of all trials of the last four blocks of that session.

#### Variability

For each speed session, we calculated the variability in gains following rewarded and following non-rewarded trials. We took a simple measure of variability as the median absolute trial-to-trial change in gain following rewarded and following non-rewarded trials based on triplets of trials (van Mastrigt et al., 2021). We assume that sensorimotor noise is represented by the variability following rewarded trials. We estimated sensorimotor noise and exploration using the additional-trial-to-trial-change method (van Mastrigt et al., 2021), which assumes that sensorimotor noise and exploration are two independent sources of variability of which the variances can be added to total observed variance. Exploration is estimated by subtracting gain variance estimates based on trial-to-trial changes following success (i.e. sensorimotor noise) from gain variance estimates based on trial-to-trial changes following failure. We thus estimate exploration as the additional variability following failure relative to following success.

#### Learning

Learning was calculated as the difference in median gain of the last 10 trials of two subsequent blocks, divided by the difference in target gain. A value of one thus corresponds to full learning, a value of zero indicates no learning. To obtain one learning measure per speed session, for each participant we took the mean of the blocks.

### Statistics

As the differences between speed sessions were not normally distributed, we used a non-parametric one-tailed paired-samples t-test to test whether stepping speed was higher in the fast session than in the slow session. We also did a non-parametric paired-samples t-test to test whether learning differed between speed sessions. We first did a non-parametric paired-samples t-test to test whether overall variability was higher in the fast session than in the slow session and whether we should do two more paired-samples t-tests to test for changes in trial-to-trial changes following success (sensorimotor noise) and following failure. To decide whether we could explore exploration in our analyses in a similar fashion as overall variability, we tested whether we found significant exploration in both sessions. As exploration estimates were not normally distributed, we used a one-tailed non-parametric one-sample t-test per session.

## Results

### Movement speed

We successfully manipulated stepping speed: stepping speed was higher in the fast session than in the slow session (p < 0.01, z = -5.18, W = 18, r = -0.13) (Fig 1a). The same pattern was observed for each gain block separately. Participants achieved the higher stepping speed in the fast session despite their lower percentage of correctly-timed steps in the fast session (80%) than in the slow session (40%) (Fig 1c). Higher stepping speed was achieved by actually stepping faster rather than taking longer steps in the slow session and shorter steps in the fast session: the range in step lengths was similar across speed sessions (Fig 1b).

**Fig 1.**
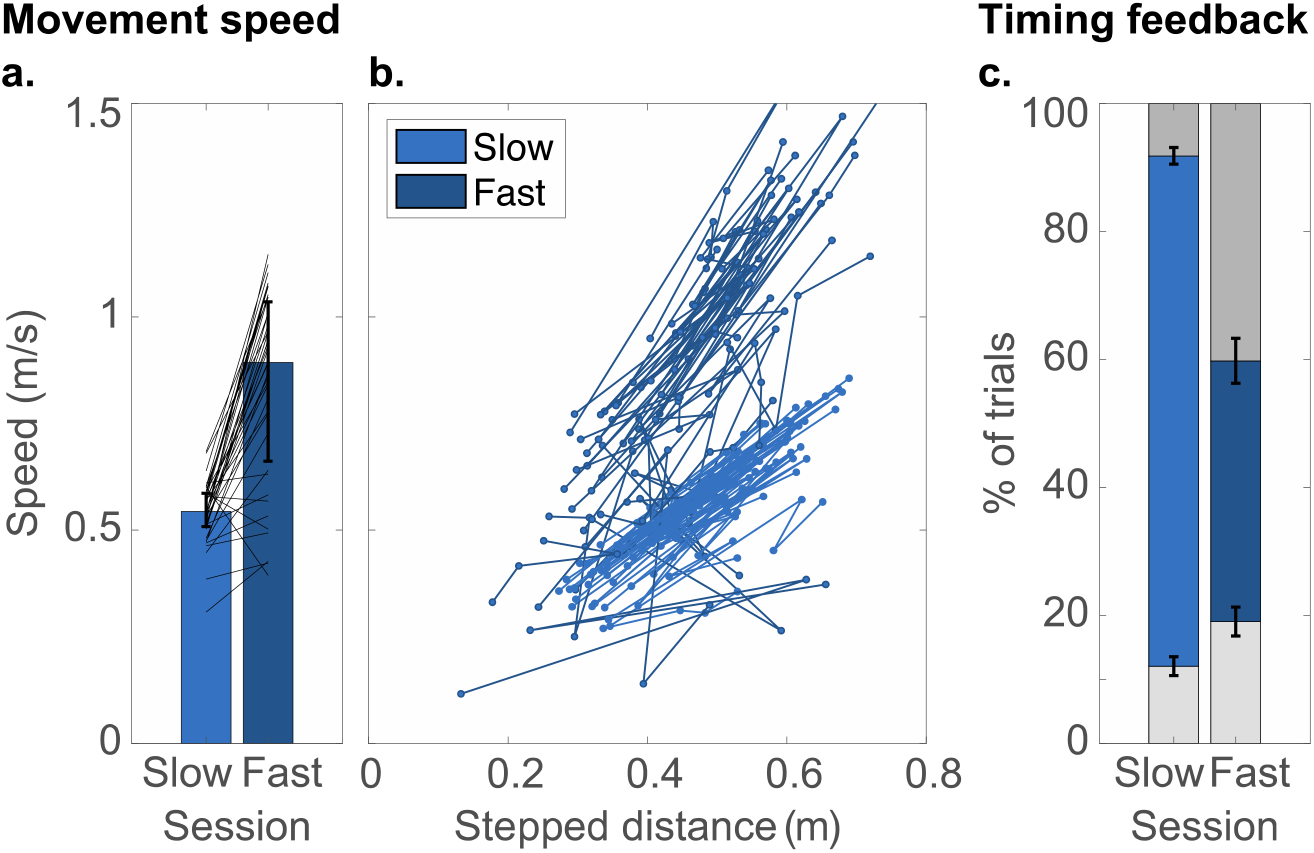
Movement speed. a. Movement speed per speed session. Lines indicate the individual participants’ data for the two speed sessions, averaged across gain blocks. The filled bars indicate the median across participants, with error bars for the interquartile range. b. Movement speed as a function of step length for individual participants’ blocks and speed sessions. The lines connect the values for the four consecutive blocks. c. Percentage of trials that were correctly timed (blue), that participants moved too fast (light grey) or too slow (dark grey).

### Learning

Participants could use the binary reward feedback to adjust their gains within gain blocks (Fig 2a). They could do so in both speed sessions (Fig 2b) and showed higher final gains in the blocks with higher target gains. On average, participants showed low to moderate learning in both the slow and fast session. We found no indication for a difference in learning between the speed sessions (p = 0.49, W = 389). The adaptive reward criterion, designed to get similar numbers of trials with success and failure, resulted in average success percentages of 48 ±5% and 47 ±6% in the slow and fast session, respectively.

**Fig 2.**
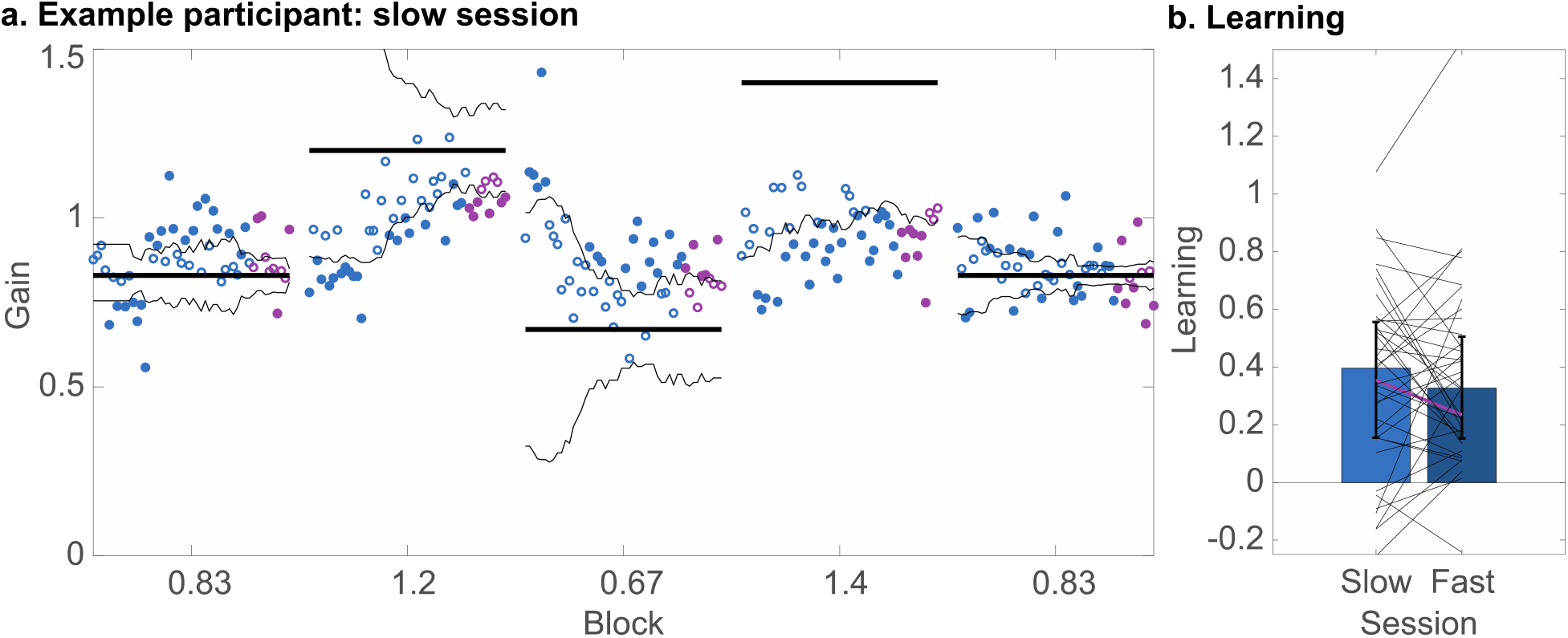
Learning. a. Gain data for an example participant in the slow session. Open circles indicate rewarded trials, filled circles indicate non-rewarded trials. Trials that were used for calculating learning are depicted in pink. The black thin lines indicate the borders of reward zone. The thick, horizontal black lines indicate the target gain. b. Learning per speed session. Error bars indicate the median and interquartile range over participants. Thin lines indicate individuals, with the example participant of panel a depicted in pink.

### Variability

We found significantly higher overall variability, as quantified by higher median absolute trial-to-trial changes in gain in the fast session than in the slow session (Fig 3a, p < 0.01, z = - 2.49, W = 211, r = -0.06). Variability following failure and following success showed the same pattern (Fig 3b): we found significantly higher median absolute trial-to-trial changes in the fast session than in the slow session, both following success (p < 0.01, z = -3.00, W = 174, r = -0.08) and following failure (p < 0.05, z = -1.78, W = 262, r = -0.05). We thus found significantly higher sensorimotor noise in the fast session (i.e. variability following-success). We estimated exploration as the additional variability following failed trials as compared to following successful trial, subtracting variance estimates following failure and following success (see Methods). Within both sessions, we found significant exploration (Slow: p < 0.05, W = 549, r = 0.06; Fast: p < 0.01, W = 594, r = 0.07), despite the fact that we observed many negative estimates and a high signal-to-noise ratio (Fig 3c). Since a negative variance cannot exist, these negative exploration estimates can be regarded as an error due to the low reliability in variance estimates following successful and non-successful trials. We found no significant difference in exploration between sessions (p = 0.27, W = 346).

**Fig 3.**
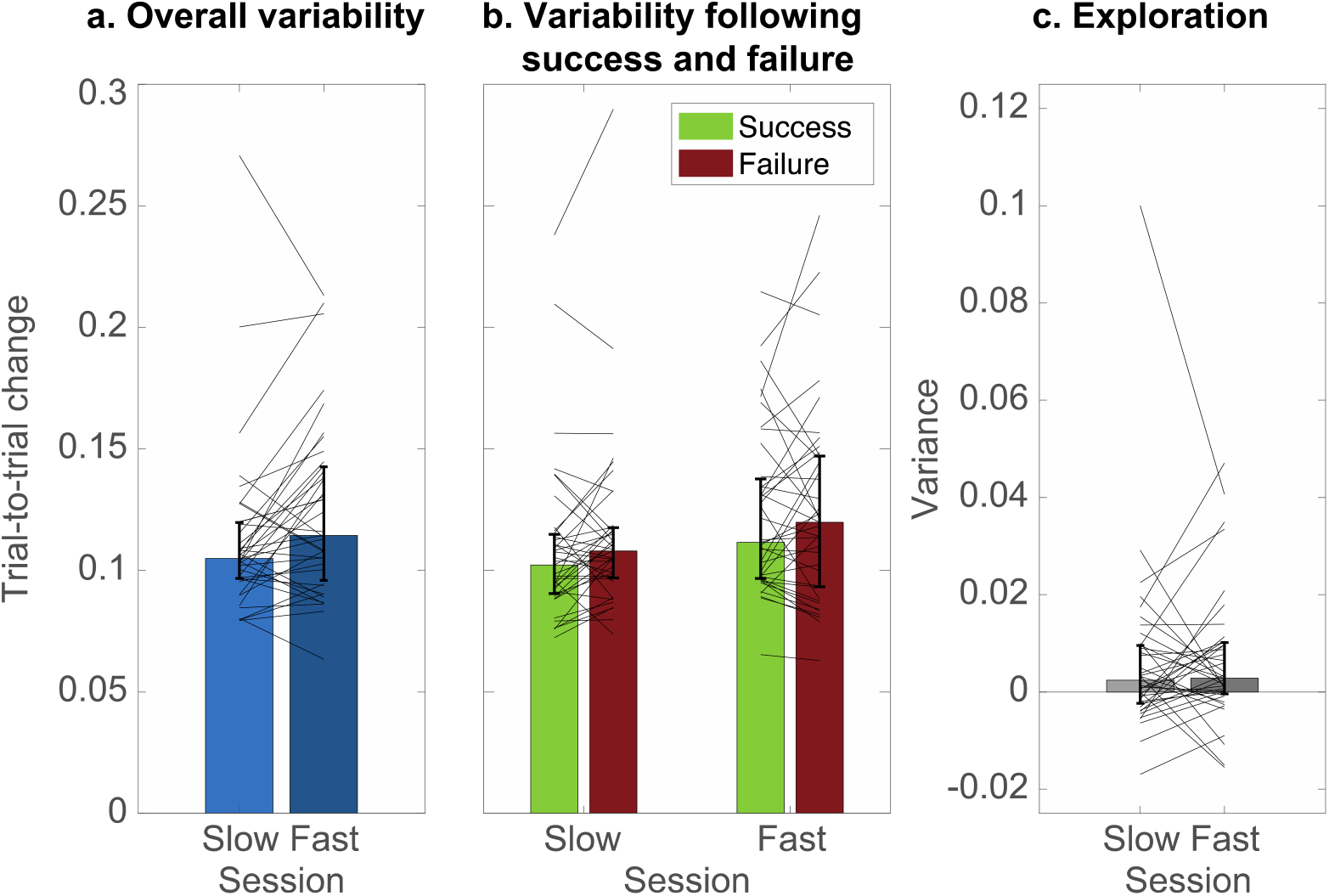
Variability. a. Variability in gain expressed as median absolute trial-to-trial change over all trials. b. Feedback-modulated variability following success and following failure, expressed as median absolute trial-to-trial change. c. Exploration, expressed as a variance. In all panels, error bars indicate the median and interquartile range over participants. Thin lines indicate individuals.

We explored the relation between variability and learning (Fig 4) to see which sources of variability were related to learning. We did so for overall variability, variability following success (i.e. sensorimotor noise), variability following failure, exploration, and the exploration as a fraction of total variability as quantified by the sum of the variance of sensorimotor noise and exploration. Since sensorimotor noise has been shown to hamper learning (Therrien et al., 2016, 2018), we had expected a negative relation between sensorimotor noise and learning (Figure 4b, g). Since exploration is generally thought of as promoting learning (Dhawale et al., 2019; Sutton & Barto, 2017; Therrien et al., 2016), we had expected a positive relation between exploration and learning (Fig 4 d, i). Since learning may depend on the balance of exploration and motor noise (Therrien et al., 2016, 2018), we had expected a positive relation between the exploration fraction of total variability and learning (Fig 4 e, j). Our five measures of variability seem however to be unrelated to the amount of learning (Fig 4, upper panels). The same is true for the difference in the respective values between speed sessions (Fig 4, lower panels).

**Fig 4.**
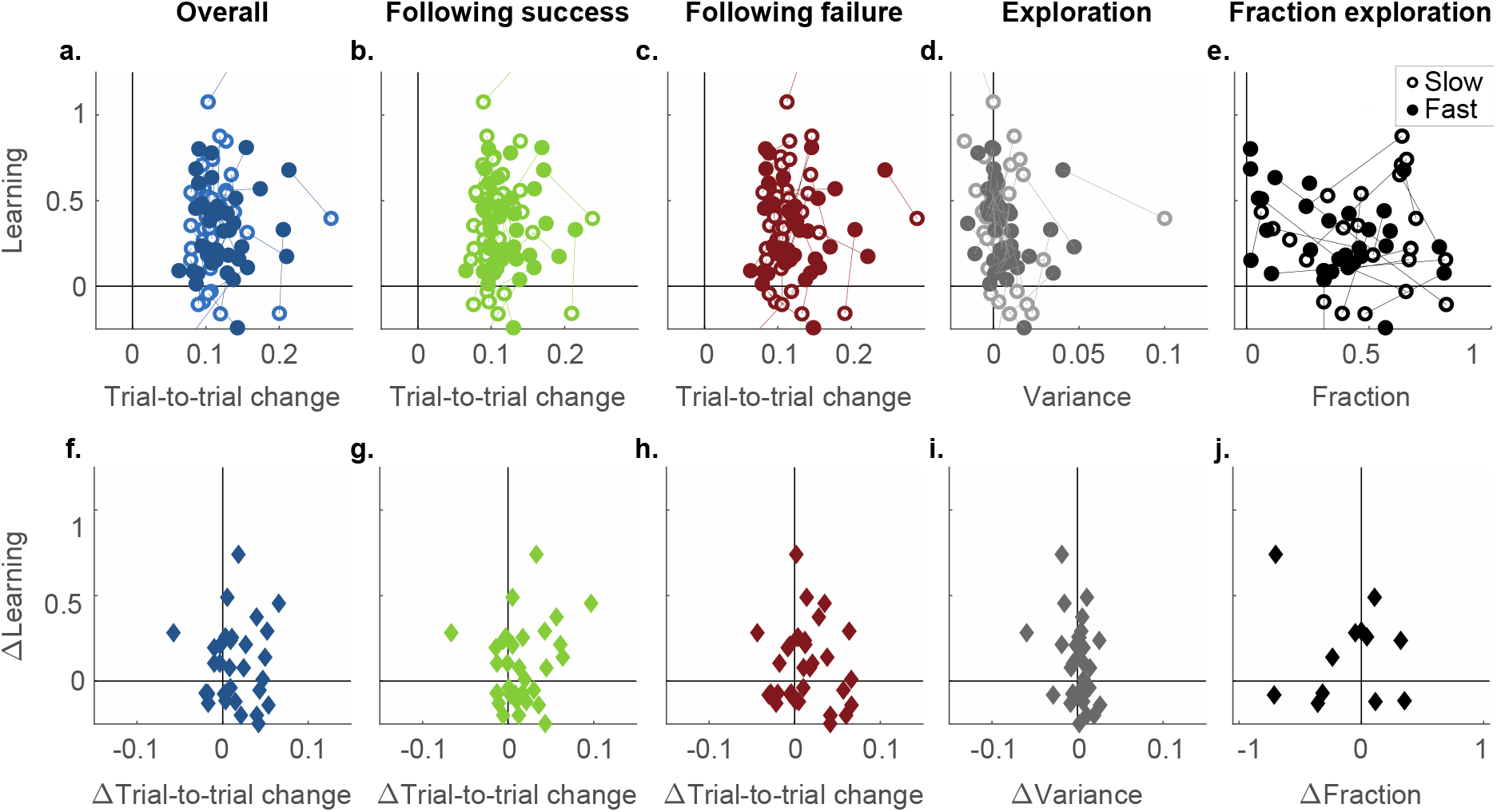
Variability and learning. a-e. Learning as a function of variability in gain. Each participant is represented by an open and closed symbol connected by a line, corresponding to the slow and fast session. f-j. Changes in learning as a function of changes in variability, from the slow to the fast session. Each participant is represented by one symbol. a, f. Overall variability. b, g. Feedback-modulated variability: median absolute trial-to-trial change following success, which is a proxy for sensorimotor noise. c, h. Feedback-modulated variability: median absolute trial-to-trial change following failure. d, i. Exploration, expressed as a variance. E, j. Fraction of exploration to total variability, expressed as exploration variance divided by the sum of the variances of sensorimotor noise and exploration. Only sessions with positive exploration estimates are depicted in panel e, and only participants who had positive exploration estimates in both sessions are depicted in panel j.

## Discussion

We aimed to answer the question whether increasing movement speed can be harnessed to promote motor learning. Participants performed a gain-learning task with binary reward feedback based on their step length. We expected that increasing movement speed would result in increases in motor variability, which in reward-based motor learning tasks may promote learning. We successfully manipulated movement speed. Gain variability increased from the slow to the fast session, both in general, following success and following failure. We found significant exploration in both speed sessions. Participants could learn the task, but we found no effect of movement speed on the amount of learning, nor on exploration.

Participants could learn to increase or decrease their gain. We found limited learning in both speed sessions. Learning was lower than in other reward-based motor learning tasks, such as center-out reaching tasks with visuomotor perturbations (Codol et al., 2018; Holland et al., 2018; Izawa & Shadmehr, 2011; Therrien et al., 2016, 2018). This is however not surprising since participants had a limited number of trials (i.e. fifty) to learn each gain as compared to the hundreds of trials of (Codol et al., 2018; Holland et al., 2018; Izawa & Shadmehr, 2011; Therrien et al., 2016, 2018). Moreover, the task is a whole-body task with multiple targets (van der Kooij & Smeets, 2018), in which a gain rather than a rotation has to be learnt. Learning may also have been limited by high levels of sensory noise (He et al., 2016) in perceiving the gap distance on screen, perceiving the distance stepped and perceiving the gain with which a participant had stepped. The limited learning may also have been the result of high levels of motor noise, which has been shown to impair reward-based learning (Therrien et al., 2018). Another explanation may be that stepping at a certain speed, by constraining movement time, may have functioned as a dual task for participants since they received feedback on both gain and timing. A dual task has been shown to reduce the amount of reward-based motor learning in a center-out reaching task (Holland et al., 2018). Lastly, the reward criterion aimed at rewarding half of the attempts may have been not informative enough to learn. All in all, we found modulation of gains and step lengths based on the feedback, indicating that participants learned.

We found reward-based adaptation of stepping gain in a task where the target distance was pseudo-randomly chosen on each trial. While it has been known that participants can adapt their reach direction to a single target based on reward-based feedback (e.g. (Cashaback et al., 2019; Izawa & Shadmehr, 2011; Therrien et al., 2016), reward-based motor learning decreases with the number of targets (van der Kooij & Smeets, 2018), and hasn’t been observed in three-dimensional tasks where participants moved to randomly chosen target positions with a constant shift of the rewarded position (van der Kooij et al., 2018; van der Kooij & Overvliet, 2016). Our results show that while reward-based adaptation might decrease with the number of target positions (van der Kooij & Smeets, 2018), reward-based adaptation is still possible with a wide range of target positions. This argues against the idea that reward-based motor learning relies on the learning of action-outcome mappings (Izawa & Shadmehr, 2011). As we find learning and exploration of gain in our stepping task, this suggests that reward-based learning plays a role in learning the visuomotor transformations as well.

Despite our manipulation of variability, we found no difference in reward-based learning between the slow and fast sessions. Movement speeds may have been too close to induce any differences in learning, but this is not likely as participants tended to be too early in the slow session and too late in the fast session (i.e. movement speed induced clearly different behaviour). Although we successfully increased gain variability from the slow to the fast session, both overall, following success and following failure, these differences may have been too small to lead to a detectable difference in learning. Alternatively, we may not have found a difference in learning because we only found evidence for an increase in sensorimotor noise with speed, and no evidence for an increase in exploration, whereas sensorimotor noise is often considered variability that cannot be learnt from (Dhawale et al., 2019; Therrien et al., 2016, 2018). Lastly, learning may have been constrained by other factors than speed or variability, making the task so difficult that it became insensitive to movement speed in both sessions.

Since we manipulated movement speed to induce variability, we explored how movement speed influenced different sources of variability and how (changes in) sensorimotor noise, exploration and their ratio were related to (changes in) learning. We found that increasing movement speed did result in higher gain variability: overall, following success and following failure. We found no evidence for a relation between any of the variability sources and learning. Our finding that total variability and sensorimotor noise (i.e. variability following success) increase with movement speed seems to be in line with speed-accuracy trade-offs reported in previous arm (Fitts, 1954) and whole-body center-of-pressure tasks (Duarte & Freitas, 2005). Since the feedback in these experiments was online error-feedback rather than external binary reward feedback, we consider their higher variability with increasing speed sensorimotor noise rather than exploration (van Mastrigt et al., 2020, 2021). As our exploration estimates were unreliable due to the low number of trials used for the ATTC method and the high levels of variability following successful trials (van Mastrigt et al., 2021), we cannot conclude whether movement speed increased exploration. If one assumes that total variability is the sum of the variances of sensorimotor noise and exploration, one could (falsely) argue that exploration did not change because sensorimotor noise did. That is however over-interpretation of non-significant results (Makin & Orban de Xivry, 2019), because we cannot be sure whether it represents a true null-result, too low statistical power to detect an effect.

In conclusion, we showed reward-based learning of a gain in a stepping task. We tested whether increasing movement speed influenced learning, and found no difference in learning between a slow and fast stepping session. Increasing movement speed led to increased levels of sensorimotor noise and total variability, but we found no relation between variability sources and learning. Neither did we find evidence for a relation between variability sources and learning. We believe the question at which speed to move when learning a motor task is a basic and interesting question, not only from our perspective of motor variability. Future experiments are needed to explore how movement speed influences different sources of variability and whether the speed-accuracy trade-off can be harnessed to promote motor learning. The task should be relatively easy to learn and have low levels of sensory noise. If movement speed is used to manipulate variability, controlling speed should not serve as a dual task alongside a learning task. Possibly, constraining movement preparation time may serve as an alternative manipulation to manipulate variability with (Sutter et al., 2021).

## Acknowledgments

The research was funded by the Nederlandse Organisatie voor Wetenschappelijk Onderzoek, Toegepaste en Technische Wetenschappen Open Technologie Programma (NWO-TTW OTP grant 15989). The funders had no role in study design, data collection and analysis, decision to publish, or preparation of the manuscript.

## Authorship

Conceptualization: Nina M. van Mastrigt, Katinka van der Kooij, Jeroen B.J. Smeets

Funding acquisition: Jeroen B. J. Smeets, Katinka van der Kooij

Investigation: Nina M. van Mastrigt.

Methodology: Nina M. van Mastrigt, Katinka van der Kooij, Jeroen B.J. Smeets

Software: Nina van Mastrigt (Matlab + psychtoolbox)

Supervision: Jeroen B. J. Smeets, Katinka van der Kooij.

Visualization: Nina M. van Mastrigt

Writing – original draft: Nina M. van Mastrigt

Writing – review & editing: Nina M. van Mastrigt, Katinka van der Kooij, Jeroen B.J. Smeets

## Data and code availability

Data and code can be accessed on the Open Science Framework (https://osf.io/x7hp9/).

## Notes

https://osf.io/x7hp9/

### Competing Interest Statement

The authors have declared no competing interest.

## References

Abram, S. J., Poggensee, K. L., Sánchez, N., Simha, S. N., Finley, J. M., Collins, S. H., & Donelan, J. M. (2022). General variability leads to specific adaptation toward optimal movement policies. Current Biology, 32, 1–11. https://doi.org/10.1016/j.cub.2022.04.015

Bernstein, N. (1967). The co-ordination and regulation of movements. (1st ed.). Pergamon Press Ltd.

Cashaback, J. G. A., Lao, C. K., Palidis, D. J., Coltman, S. K., McGregor, H. R., & Gribble, P. L. (2019). The gradient of the reinforcement landscape influences sensorimotor learning. PLoS Computational Biology, 15(3), e1006839. https://doi.org/10.1371/journal.pcbi.1006839

Codol, O., Holland, P. J., & Galea, J. M. (2018). The relationship between reinforcement and explicit control during visuomotor adaptation. Scientific Reports, 8(9121), 1–11. https://doi.org/10.1038/s41598-018-27378-1

Dhawale, A. K., Miyamoto, Y. R., Smith, M. A., & Ölveczky, B. P. (2019). Adaptive Regulation of Motor Variability. Current Biology, 29(21), 3551–3562.e7. https://doi.org/10.1016/j.cub.2019.08.052

Duarte, M., & Freitas, S. M. S. F. (2005). Speed-accuracy trade-off in voluntary postural movements. Motor Control, 9(2), 180–196. https://doi.org/10.1123/mcj.9.2.180

Fitts, P. M. (1954). The information capacity of the human motor system in controlling the amplitude of movement. Journal of Experimental Psychology, 47(6), 381–391.

Harris, C. M., & Wolpert, D. M. (1998). Signal-dependent noise determines motor learning. Nature, 394(August), 780–784. https://doi.org/10.1038/nature01718.1.

He, K., Liang, Y., Abdollahi, F., Fisher Bittmann, M., Kording, K., & Wei, K. (2016). The Statistical Determinants of the Speed of Motor Learning. PLoS Computational Biology, 12(9), 1–20. https://doi.org/10.1371/journal.pcbi.1005023

Herzfeld, D. J., & Shadmehr, R. (2014). Motor variability is not noise, but grist for the learning mill. Nature Neuroscience, 17(2), 149–150. https://doi.org/10.1038/nn.3633

Holland, P., Codol, O., & Galea, J. M. (2018). Contribution of explicit processes to reinforcement-based motor learning. Journal of Neurophysiology, 119(6), 2241–2255. https://doi.org/10.1152/jn.00901.2017

Izawa, J., & Shadmehr, R. (2011). Learning from sensory and reward prediction errors during motor adaptation. PLoS Computational Biology, 7(3), e1002012. https://doi.org/10.1371/journal.pcbi.1002012

Makin, T. R., & Orban de Xivry, J.-J. (2019). Ten common statistical mistakes to watch out for when writing or reviewing a manuscript. ELife, 8, 1–13. https://doi.org/10.7554/eLife.48175

Mathôt, S., Schreij, D., & Theeuwes, J. (2012). OpenSesame: An open-source, graphical experiment builder for the social sciences. Behavior Research Methods, 44(2), 314–324. https://doi.org/10.3758/s13428-011-0168-7

Pekny, S. E., Izawa, J., & Shadmehr, R. (2015). Reward-Dependent Modulation of Movement Variability. Journal of Neuroscience, 35(9), 4015–4024. https://doi.org/10.1523/JNEUROSCI.3244-14.2015

Sutter, K., Wijdenes, L. O., Van Beers, R. J., & Medendorp, W. P. (2021). Movement preparation time determines movement variability. Journal of Neurophysiology, 125(6), 2375–2383. https://doi.org/10.1152/jn.00087.2020

Sutton, R. S., & Barto, A. G. (2017). Reinforcement learning: an introduction (2nd ed.). MIT Press.

Therrien, A. S., Wolpert, D. M., & Bastian, A. J. (2016). Effective Reinforcement learning following cerebellar damage requires a balance between exploration and motor noise. Brain, 139(1), 101–114. https://doi.org/10.1093/brain/awv329

Therrien, A. S., Wolpert, D. M., & Bastian, A. J. (2018). Increasing Motor Noise Impairs Reinforcement Learning in Healthy Individuals. Eneuro, 5(June), ENEURO.0050-18.2018. https://doi.org/10.1523/ENEURO.0050-18.2018

Tumer, E. C., & Brainard, M. S. (2007). Performance variability enables adaptive plasticity of “crystallized” adult birdsong. Nature, 450(7173), 1240–1244. https://doi.org/10.1038/nature06390

Uehara, S., Mawase, F., Therrien, A. S., Cherry-Allen, K. M., & Celnik, P. A. (2019). Interactions between motor exploration and reinforcement learning. Journal of Neurophysiology, 122, 797–808. https://doi.org/10.1152/jn.00390.2018

van Beers, R. J. (2009). Motor Learning Is Optimally Tuned to the Properties of Motor Noise. Neuron, 63(3), 406–417. https://doi.org/10.1016/j.neuron.2009.06.025

van der Kooij, K., & Overvliet, K. E. (2016). Rewarding imperfect motor performance reduces adaptive changes. Experimental Brain Research, 234(6), 1441–1450. https://doi.org/10.1007/s00221-015-4540-1

van der Kooij, K., & Smeets, J. B. J. (2018). Reward-Based Motor Adaptation Can Generalize Across Actions. Journal of Experimental Psychology: Learning Memory and Cognition, 0(999), 71–81. https://doi.org/10.1037/xlm0000573

van der Kooij, K., van Mastrigt, N. M., Crowe, E. M., & Smeets, J. B. J. (2021). Learning a reach trajectory based on binary reward feedback. Scientific Reports, 11(1), 1–15. https://doi.org/10.1038/s41598-020-80155-x

van der Kooij, K., Wijdenes, L. O., Rigterink, T., Overvliet, K. E., & Smeets, J. B. J. (2018). Reward abundance interferes with error-based learning in a visuomotor adaptation task. PLoS ONE, 13(3), e0193002. https://doi.org/10.1371/journal.pone.0193002

van der Vliet, R., Frens, M. A., de Vreede, L., Jonker, Z. D., Ribbers, G. M., Selles, R. W., van der Geest, J. N., & Donchin, O. (2018). Individual Differences in Motor Noise and Adaptation Rate Are Optimally Related. Eneuro, 5(4), ENEURO.0170-18.2018. https://doi.org/10.1523/ENEURO.0170-18.2018

van Mastrigt, N. M., Smeets, J. B. J., & van der Kooij, K. (2020). Quantifying exploration in reward-based motor learning. Plos ONE, 15(4), e0226789. https://doi.org/10.1371/journal.pone.0226789

van Mastrigt, N. M., van der Kooij, K., & Smeets, J. B. J. (2021). Pitfalls in quantifying exploration in reward-based motor learning and how to avoid them. Biological Cybernetics, 115(4), 365–382. https://doi.org/10.1007/s00422-021-00884-8

Wu, H. G., Miyamoto, Y. R., Castro, L. N. G., Ölveczky, B. P., & Smith, M. A. (2014). Temporal structure of motor variability is dynamically regulated and predicts motor learning ability. Nature Neuroscience, 17(2), 312–321. https://doi.org/10.1038/nn.3616

